# ngsReports: An R Package for managing FastQC reports and other NGS related log files

**DOI:** 10.1101/313148

**Authors:** Christopher M. Ward, Hein To, Stephen M Pederson

## Abstract

**Motivation:** High throughput next generation sequencing (NGS) has become exceedingly cheap facilitating studies to be undertaken containing large sample numbers. Quality control (QC) is an essential stage during analytic pipelines and can be found in the outputs of popular bioinformatics tools such as FastQC and Picard. Although these tools provide considerable power when carrying out QC, large sample numbers can make identification of systemic bias a challenge.

**Results:** We present ngsReports, an R package designed for the management and visualization of NGS reports from within an R environment. The available methods allow direct import into R of FastQC output as well as that from aligners such as HISAT2, STAR and Bowtie2. Visualization can be carried out across many samples using heatmaps rendered using *ggplot2* and *plotly*. Moreover, these can be displayed in an interactive shiny app or a HTML report. We also provide methods to assess observed GC content in an organism dependent manner for both transcriptomic and genomic datasets. Importantly, hierarchical clustering can be carried out on heatmaps with large sample sizes to quickly identify outliers and batch effects.

**Availability and Implementation:** ngsReports is available at https://github.com/UofABioinformaticsHub/ngsReports.

## Introduction

The Next Generation Sequencing (NGS) boom of genetics has provided researchers unparalleled resources to answer fundamental questions in population and medical genetics. The cost of sequencing has been decreasing steadily since the advent of second generation sequencing, leading to an increase in data being generated.

Quality control (QC) is arguably the most important stage in bioinformatic analysis and pipeline optimization. One of the most heavily used tools to aid in the identification of systematic and user-induced bias for NGS data is FASTQC (https://www.bioinformatics.babraham.ac.uk/projects/fastqc/), which provides a series of detailed statistics and plots for an individual fastq or bam file. In addition, many tools, such as BowTie2 (Langmead and Salzberg, 2012), HISAT2 (Kim, et al., 2015) and STAR (Dobin, et al., 2013), provide useful summaries and statistics which can be saved to disk. However, collating and interpreting quality control logs across multiple files can be tedious and prone to human error as most tools output logs on a per-sample basis. The decrease in cost and increase in popularity of NGS-based experimental designs further contributes to the growing need of software to collate, manipulate and visualize QC logs and reports.

Few software tools are currently available to tackle this problem, with the notable exception of MultiQC (Ewels, et al., 2016). Although useful at investigating multiple samples for errors, MultQC functions well as a standalone tool, however summaries and reports are often required for researchers working within an R environment. Therefore we present ngsReports, the first R package to incorporate NGS log files and quality control into the R programming language. We seek to provide a platform to easily and accurately compare NGS logs to reduce human error in the identification of outliers and batch effects and to simplify pipeline optimization.

## Methods

### Data analysis

To aid in access, manipulation, and storage of data we created novel S4 classes for FastQC reports (FastqcFile & FastqcData), whilst multiple reports are collated into list-like extensions (FastqcFileList & FastqcDataList). Collated S4 objects can be subset using standard list nomenclature and passed directly into plotting functions. Data for each FastQC module is accessible by passing the FastqcDataList into individual functions (eg. Per_base_sequence_quality). Data can then be freely manipulated using R packages such as *dplyr* (Wickham, et al., 2017) and *reshape2* (Wickham, 2007). We also provide functions to parse and import log files of popular bioinformatics tools: Bowtie (Langmead, et al., 2009) & Bowtie2 (Langmead and Salzberg, 2012), HISAT2 (Kim, et al., 2015), Picard:Mark Duplicates (http://broadinstitute.github.io/picard/) and STAR (Dobin, et al., 2013).

### Visualisation

Each module of the FastQC report is contained within its own S4-method and can be rendered using either *ggplot2* (Wickham, 2016) or *plotly* (https://github.com/ropensci/plotly). Plots for each module of the FastQC report can be generated for single or multiple files in either line or heatmap format (Fig. 1) using functions provided. Plots rendered using *ggplot2* remain compatible with the addition of themes, annotations and geom’s using standard *ggplot2* nomenclature. Plots can be made interactive using the *usePlotly* argument; this also displays a sidebar denoting the Pass/Warning/Fail flag for that module in the FastQC report (Fig. 1). Hierarchical clustering in heatmaps and barplots is also supported to allow for fast identification of outlier libraries and batch effects (Fig. 1).

**Figure 1:**
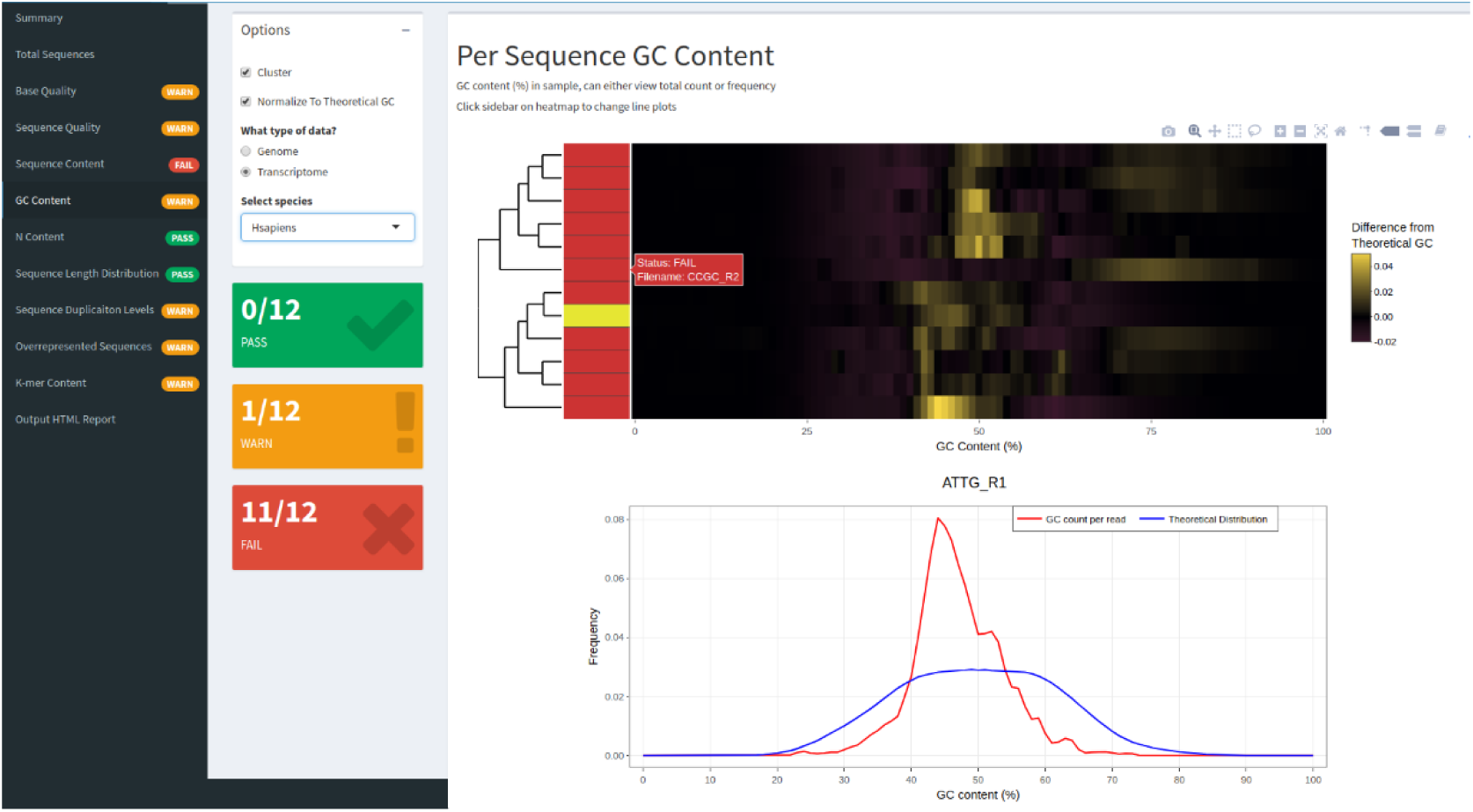
Visualization methods in ngsReports. The FASTQC shiny app, provides interactive and customizable plots for each FastQC module (tabs). FastQC reports can be input directly into the shiny app, with the option to output a HTML report when finished. Here we show a clustered heatmap with dendrogram displaying the difference between the observed GC content and the theoretical GC for *Homo sapien* genome short reads across all reads. Sidebar of the plot shows the PASS/WARN/FAIL status attributed to the “GC content” module by FastQC. **B)** The line plot for a single individual, ‘ATTG_R1,’ this can be carried out on-click in the shiny app to provide more detailed information about an individual sample.

Observed GC content in libraries can be visualised using one of two methods. Total observed GC content (Supplementary Fig. 1) or after subtractive normalisation using the theoretical GC content for a closely related organism of interest (Fig. 1). Theoretical GC content has been prepared and supplied for many model organisms (https://github.com/mikelove/fastqcTheoreticalGC) for both transcriptomic and genomic datasets; this allows biases in GC content to be identified in an organism and -omic dependent manner.

For easy integration with data processing pipelines, an interactive FastQC summary HTML report can also be produced using a custom R markdown template, with a default template provided with the package. This gathers summary tables and FastQC module plots into a single file for ease of analysis.

### Shiny App

In addition to the HTML report, we also provide a GUI written using *shiny*. This provides an interactive user interface to view and customize plots for each module within the FastQC report (Fig. 1). Furthermore, the shiny app displays arguments for clustering, zooming, data type and organism selection and on-click inspection of individual line plots for libraries (Fig. 1).

## Conclusion

Costs involved in library preparation and sequencing for NGS applications are decreasing, allowing for much larger studies to be undertaken. The functional programming capabilities of the R programming language provide an ideal environment for data manipulation and analysis. The methods provided in ngsReports constitute a powerful tool for generic and bespoke aggregation, analysis and visualization of NGS quality control and log data.

## References

Dobin, A., et al. (2013) STAR: ultrafast universal RNA-seq aligner, Bioinformatics, 29, 15-21.

Ewels, P., et al. (2016) MultiQC: summarize analysis results for multiple tools and samples in a single report, Bioinformatics, 32, 3047-3048.

Kim, D., Langmead, B. and Salzberg, S.L. (2015) HISAT: a fast spliced aligner with low memory requirements, Nature Methods, 12, 357.

Langmead, B. and Salzberg, S.L. (2012) Fast gapped-read alignment with Bowtie 2, Nature methods, 9, 357-359.

Langmead, B., et al. (2009) Ultrafast and memory-efficient alignment of short DNA sequences to the human genome, Genome Biology, 10, R25.

Wickham, H. (2007) Reshaping Data with the reshape Package, 2007, 21, 20.

Wickham, H. (2016) ggplot2: elegant graphics for data analysis. Springer.

Wickham, H., et al. (2017) dplyr: A Grammar of Data Manipulation, https://CRAN.R-project.org/package=dplyr.

